# Hyperspectral environmental illumination maps: Characterizing directional spectral variation in natural environments

**DOI:** 10.1101/660290

**Authors:** Takuma Morimoto, Sho Kishigami, João M.M. Linhares, Sérgio M.C. Nascimento, Hannah E. Smithson

## Abstract

Objects placed in real-world scenes receive incident light from every direction, and the spectral content of this light may vary from one direction to another. In computer graphics, environmental illumination is approximated using maps that specify illumination at a point as a function of incident angle. However, to-date, existing public databases of environmental illumination maps specify only three colour channels (RGB). We have captured a new set of 12 environmental illumination maps (eight outdoor scenes; four indoor scenes) using a hyperspectral imaging system with 33 spectral channels. The data reveal a striking directional variation of spectral distribution of lighting in natural environments. We discuss limitations of using daylight models to describe natural environmental illumination.

## 1. Introduction

The vast majority of objects do not themselves emit light and are therefore only visible by virtue of the incident light they reflect. Characterizing the properties of natural lighting is therefore important, as it allows us to comprehensively understand the sets of physical inputs that visual sensors receive in everyday life. The variation of manufactured indoor lighting is relatively accessible (e.g. [1]), but natural daylights require systematic investigation. In past studies, vigorous efforts have been made to characterize the spectral variation of illumination that can occur in a natural outdoor environment. The most widely known is the CIE daylight model [2], which accounts for the major variations in the spectral content of natural daylight using only three basis functions. Other studies show similar statistical trends where the chromaticites of natural illuminants cluster closely around the daylight locus [3]. It is also well understood how the spectrum of daylight can change over time and atmospheric conditions [4]. In addition, recent work has expanded these observations to twilight [5]. The focus of these studies has been to characterize the dominant illumination in a scene. Such measurements have significantly advanced our understanding of the human visual system, especially of mechanisms underlying the perception of objects’ surface properties, such as colour and shape. For example, in order to extract information about surface colour, the visual system might exploit the constraint that the illumination in a natural environment is likely daylight [6].

However, the fact that the spectral content of local illumination varies as a function of position in a complex scene suggests a significant influence of nearby objects in reflecting light and thereby modifying its spectral content away from the daylight locus [7]. Work has also been undertaken to characterize the directional variation in lighting, with particular emphasis on the directional distribution of intensity [8, 9]. The so-called “light from above” heuristic has received a lot of attention as a candidate Bayesian prior to interpret the shape of objects from the variation in shading across their surface [10, 11]. More recent work also suggests that visual behaviours might even be sensitive to the complex spatial structure of natural lighting [8, 12].

The present study measured the directional variation in spectral content of environmental illumination. Natural illumination mostly originates from two sources: (i) direct emission from a light source and (ii) secondary (or higher-order) reflection from other surfaces in the scene. The spectral shape of light in the secondary reflection is modified by the surface spectral reflectance of the objects. If we imagine an object placed at a specific point in a scene, the spectral content of illumination hitting the object may vary significantly as a function of incident angle. In computer graphics this directional dependency of illumination is approximated using an environmental illumination map [13], which, at each pixel in the map, stores the intensity of a light coming from a particular direction towards a single point in the scene (e.g. an object or an eye). Environmental illumination maps have proved to be important in achieving realistic appearance when rendering computer-graphics generated images of objects. However, existing databases of environmental illumination maps store only three-channel (RGB) measurements of light, and their use in perceptual science has been limited to characterizing intensity variation [14]. Consequently, the spectral variation in directional lighting is largely unexplored.

In this paper, we report new hyperspectral measurements of environmental illumination, which specify the spectral content of incident light from every direction at a point in a scene. The analysis of the collected datasets revealed that the colours of illuminations are not limited to fall along the daylight locus, and instead can deviate significantly from this locus. It was also clearly shown that there is a striking directional variability in illumination in natural scenes. This variation was observed mainly as a function of elevation angle rather than azimuth angle. For outside scenes, lights from above tend to be intense and dominated by short wavelengths (typically evoking a bluish percept), whereas lights from below are likely to have one log unit lower intensity and lower colour temperature. It was also found that indoor scenes have relatively uniform illumination across angle.

## 2. Method

### 2.1. Hyperspectral imaging system

The hyperspectral imaging system consists of a low-noise Peltier-cooled digital camera that has a spatial resolution of 1344 × 1024 pixels (Hamamatsu, model C4742-95-12ER, Hamamatsu Photonics K.K., Japan) equipped with a tunable liquid-crystal filter (VariSpec, model VS-VIS2-10-HC-35-SQ, Cambridge Research & Instrumentation, Inc., MA). Focal length was adjusted before each measurement and aperture was set to F16 to achieve a large depth of focus. The intensity resolution was 12-bits per pixel. Spectral measurements ranged from 400 nm to 720 nm in 10 nm steps. Each wavelength acquisition required a different exposure time to integrate a sufficient amount of light. Thus, before each image acquisition, we determined the exposure time separately for each wavelength using an automatic procedure so that the maximum pixel value in the acquired image fits within 85±5 percent of the upper limit of dynamic range, per individual wavelength. Immediately after an image was acquired, we measured the reflected light from a flat gray reflectance surface corresponding to Munsell N7, oriented perpendicular to the imaging axis and placed immediately adjacent to the mirror sphere, using a spectrophotomer with traceable calibration (PR-650, Photo Research Inc., Chatsworth, CA) for the purpose of spectral radiance calibration. A more detailed specification of the system can be found elsewhere [15, 16].

### 2.2. Measurement

All measurements were performed in Braga, Portugal. We collected data in eight outdoor scenes and four indoor scenes at Museu Nogueira da Silva and at the University of Minho, Gualtar Campus. We used a mirror sphere (AISI 52100 chrome steel ball, Simply Bearings Ltd, Lancashire, UK) of 3-inches diameter to enable imaging of environmental illumination over the full 4-pi radians of solid angle. Before starting a measurement, we set up the imaging system as shown in Figure 1 (a). The mirror sphere was placed on top of an optical support attached to a tripod. The height between the middle of the sphere and the ground was fixed at 158 cm. The zoom of the lens was maximized to allow the greatest physical distance between the sphere and the imaging apparatus, thereby minimizing the size of the image of the apparatus reflected in the mirror sphere. The hyperspectral camera was positioned 89 cm from the sphere so that the image of the mirror sphere fits just within upper and lower limit of the overall image, as shown in panel (b) of Figure 1. The spatial resolution of each sphere region was a 1024 × 1024. Before each acquisition, the camera alignment was adjusted to horizontal using a spirit level. To avoid direct sunlight, all outside measurements were performed in shadow. The mirror sphere reflects lights from a wide incident angle towards camera, but there are two issues if we try to construct an environmental illumination maps from only a single image of the sphere. Firstly, the scene behind the sphere is a “blind spot” for the sphere. Secondly, the center of the sphere has the reflection of the imaging apparatus itself. To overcome these issues, four images of the sphere were taken, separated by 90 degrees, as shown in panel (c) of Figure 1. The exposure time of each wavelength acquisition was determined by the aforementioned automatic procedure before the measurement and it was kept constant for all angles. Finally, a dark field image was acquired with the same exposure time for calibration purposes using the hyperspectral camera with the lens cap on. Images were processed, transformed to an equirectangular image, like a world map, and stitched together to create a full panorama image as described in the next subsection.

**Fig. 1.**
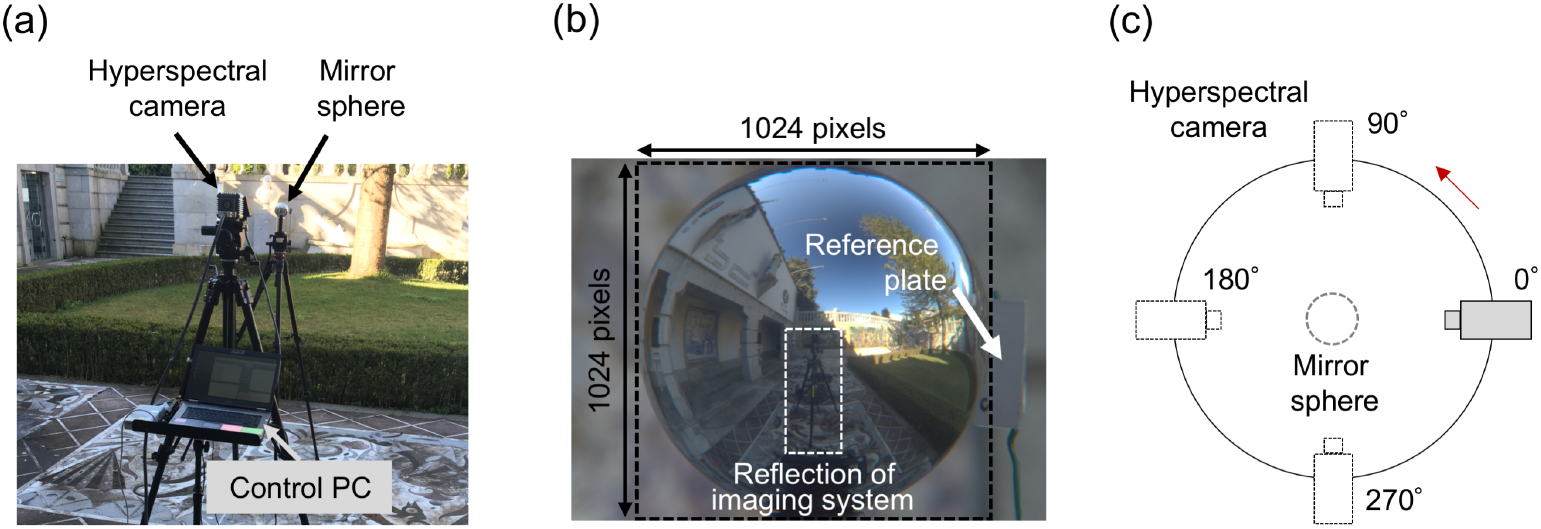
How to measure environmental illumination. (a) Set-up of the hyperspectral imaging system. The control PC was placed behind the camera so that we can erase it later along with the tripod. (b) Mirror sphere with a reference plate for calibration purposes. The spatial resolution of the sphere region was 1024 × 1024. (c) Schematic illustration of the measurement geometry. To cover the “blind-spot” of the sphere and to erase the reflection of the imaging apparatus, measurements were taken from four different angles.

### 2.3. Image Processing

Figure 2 shows the image-processing pipeline to produce an environmental illumination map from a set of mirror-sphere images. First, (i) a raw image *I*_*Raw*_(*x*,*y*;*λ*) of the sphere was corrected for dark noise, and spatial in-homogeneity. Let the dark field image be *I*_*DF*_(*x*, *y*;*λ*) and the flat field image be *I*_*FF*_(*x*,*y*;*λ*). The corrected image *I*_1_(*x*,*y*;*λ*) is given by equation (1).

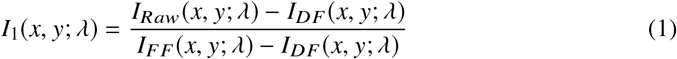

**Fig. 2.**
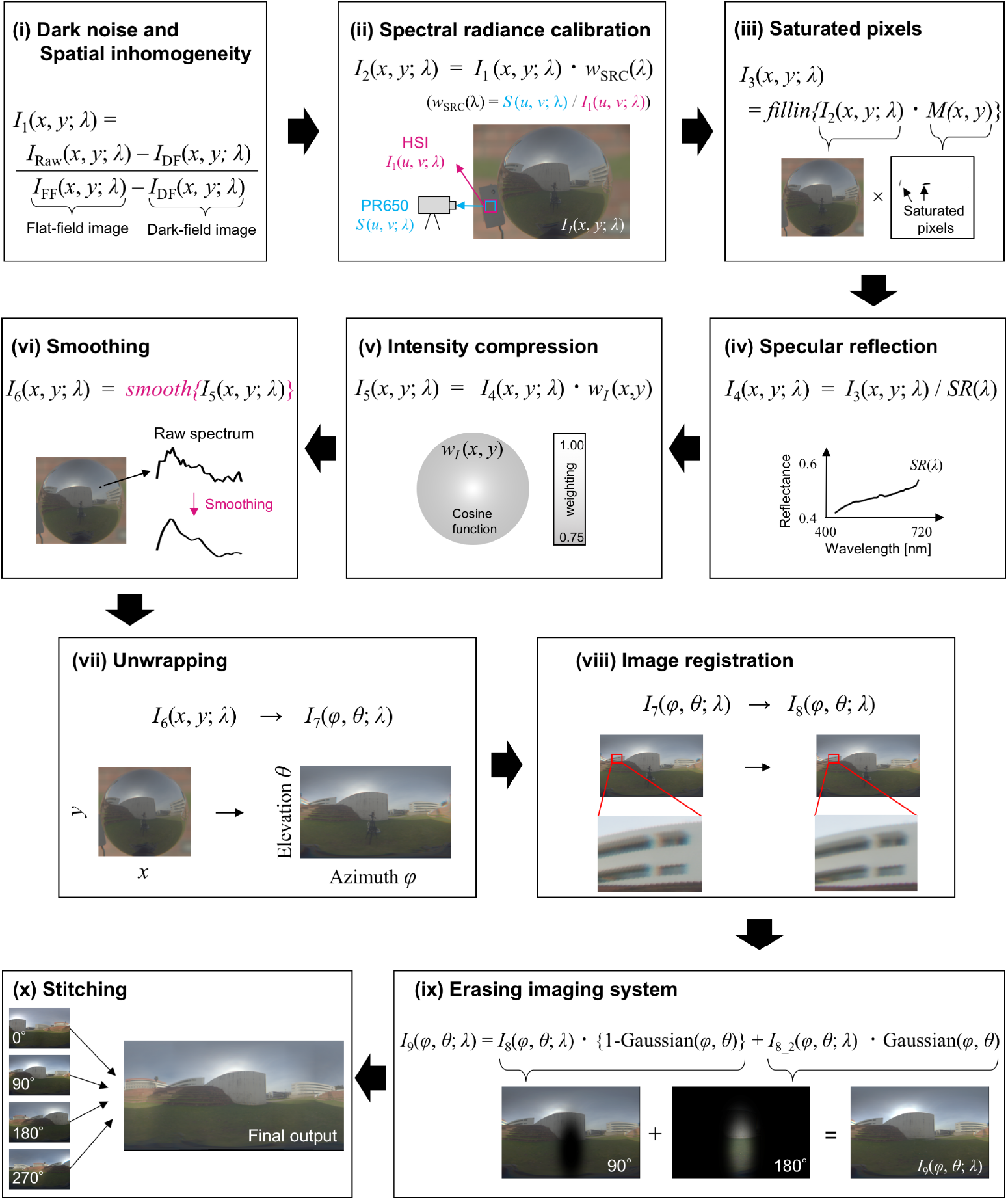
The image-processing pipeline. (i) A dark field image was measured with the imaging system with the lens cap on, and was subtracted from a raw image. Also, spatial inhomogeneites of the imaging system were corrected. (ii) Spectral radiance calibration was performed by equating the measurement from the hyperspectral imaging system to the reference measurement from the PR650 spectrophotometer. (iii) Any saturated pixels were removed, and filled-in using linear interpolation of horizontal nearest pixels. (iv) Specular reflection of the mirror sphere was measured by spectrophotometer and corrected. (v) The edge of the sphere receives and integrates lights from a wider region of the surrounding scene. This intensity compression was corrected using a cosine function of radius. (vi) Smoothing based on locally-weighted quadratic regression was performed to each pixel independently. (vii) The image coordinates were transformed to equirectangular projection. (viii) Image registration was performed to minimize the effects of chromatic aberration. (ix) The tripod was erased from each sphere image using an image from another angle. (x) Finally, four images taken from different angles were stitched together based on an automatic feature detection algorithm to create a full-panorama hyperspectral environmental illumination map.

At this stage, the image has arbitrary units. Thus, secondly, (ii) we further calibrated the image so that each pixel has units of spectral radiance (W ⋅ sr/m^2^/nm). Our approach here is to find a global scaling function that equates the spectrum at the gray reference plate recorded by the hyperspectral image to the spectrum that was independently measured in spectral radiance by a spectrophotomter PR650. Let (*u*,*v*) be the region on the gray reference plate, a spectrum reflected back from this surface measured by PR650 be *S*(*u*,*v*;*λ*) and spectra at the corresponding region in the hyperspectral image be *I*_1_(*u*,*v*;*λ*). The desired scaling function *w*_*SRC*_(*λ*) is given by equation (2).

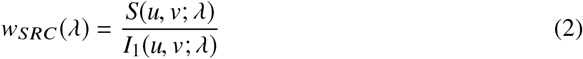

Then, we scaled whole hyperspectral image using this factor as shown in equation(3).

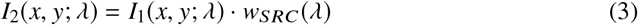

Thirdly, as shown in equation (4), (iii) any saturated pixels were removed from the image using a binary mask *M*(*x*,*y*;*λ*), where saturated and non-saturated pixels have zero and one at every wavelength, respectively. After the calculation, the resultant blank pixels were filled-in using linear interpolation of horizontal nearest pixels.

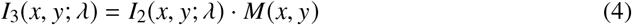

Fourth, (iv) the wavelength-dependent reflection function of the mirror sphere *SR*(*λ*) measured by spectrophotometer (Shimadzu, UV-3600 Plus UV-VIS-NIR) was corrected by equation (5).

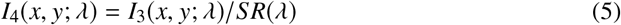

Fifth, (v) since regions close to the edge of the mirror sphere receive (and integrate) lights from a wider range of angles in the scene, this intensity compression was corrected by a cosine function of radius *w*_*I*_(*x*,*y*) as in equation (6).

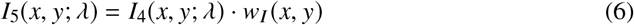

Sixth, (vi) for each pixel independently, we smoothed the spectrum based on a locally weighted quadratic regression with an optimized span of 25% using a leave-one-out cross-validation method [17] as in equation (7).

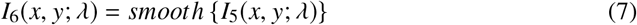

Seventh, (vii) each sphere image was converted to equirectangular coordinates as in equation (8).

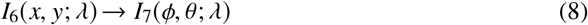

Eighth, (viii) image registration was performed using the image from the 560 nm channel as a reference to correct for chromatic aberration [18] as in equation (9).

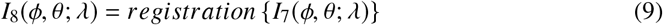

Ninth, (ix) we removed the reflection of the imaging system using an image from another angle *I*_8_2_(*φ,θ;λ*) and a gaussian filter *Gaussian*(*φ,θ*) as in equation (10).

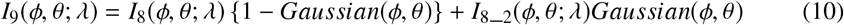

Finally, tenth, (x) four sphere images were stitched together based on the SIFT automatic feature detection algorithm [19] to produce the four-pi steradian full panorama environmental illumination map. Parts of the methods described here are analogous to those described in a recent tutorial paper [20].

## 3. Results and Discussion

Figure 3 shows the sRGB representation of the collected hyperspectral environmental illumination maps for eight outdoor scenes and four indoor scenes (panels (a) and (b) respectively). The panels (c) and (d) show the corresponding chromatic distributions of these maps expressed in the MacLeod-Boynton chromaticity diagram [21]. The horizontal axis is L/(L+M) that expresses the ratio of signals in the long and middle-wavelength sensitive cones while the vertical axis indicates S/(L+M) that expresses the signal in the short-wavelength cones in relation to a combination of long and middle-wavelength sensitive cone signals. We used the Stockman and Sharpe cone fundamentals [22] to calculate LMS cone excitations. It is clear that the chromaticities of the illumination spectra distribute around the daylight locus, but that they also deviate substantially from the locus. This was consistent for both outdoor and indoor scenes. Thus, natural lighting is not only limited to the daylight locus, but instead includes a wider range of chromaticities. The white cross indicates the chromaticity of the mean spectrum across all pixels. For outdoor scenes, it has generally lower L/(L+M) and higher S/(L+M) in relation to the chromaticity of equal energy white shown by the intersection between gray dotted lines. For indoor scenes, the mean chromaticity is slightly shifted toward lower values of S/(L+M).

**Fig. 3.**
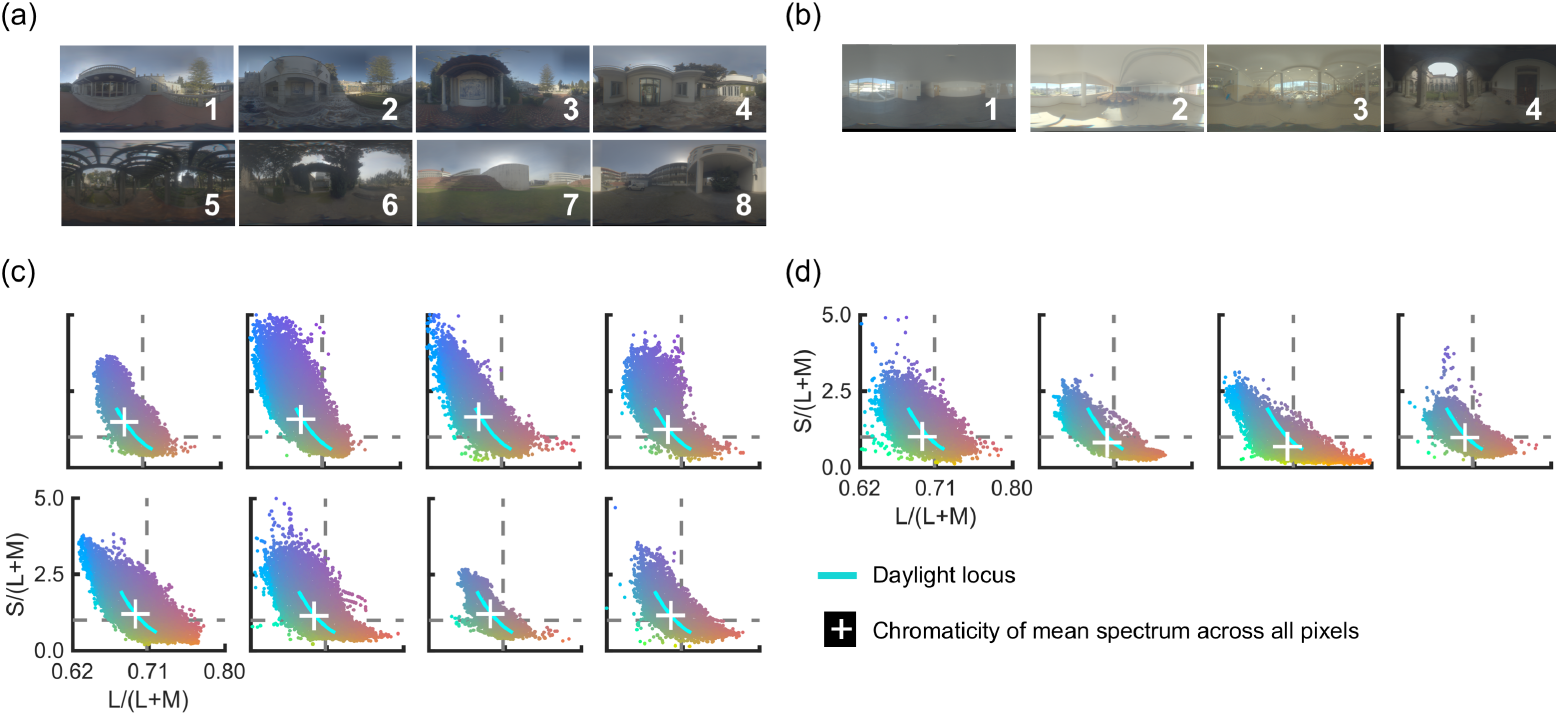
sRGB images of hyperspectral environmental illumination at (a) 8 outdoor scenes and (b) 4 indoor scenes. (c) and (d) show the chromatic distributions in the MacLeod-Boynton chromaticity diagram for each scene. The cyan line indicates the daylight locus. The gray dotted lines show the chromaticity of equal energy white.

More importantly, Figure 4 shows that there is a considerable variation in spectral shape and intensity level depending on incident angle. Each plot on the right-hand panel indicates the mean spectrum across the corresponding rectangle depicted in the left-hand panel. The background colour of each plot indicates the log luminance level. For the campus scene (a), lights from above (theta >0) tend to have high intensities in the order of thousands cd/m^2^ and high energy around the short-wavelength region. In contrast, lights reflected back from the ground and other objects, which are dominated by secondary (or higher-order) reflections, tend to have one log unit lower luminance and lower correlated colour temperature. The variation across azimuth angles (phi) is much smaller. These data clearly showed that the “light from above” heuristic does not hold universally in natural environments because of the presence of secondary reflections. In summary, the obtained data-sets revealed the large directional spectral variation in real-world visual environments.

**Fig. 4.**
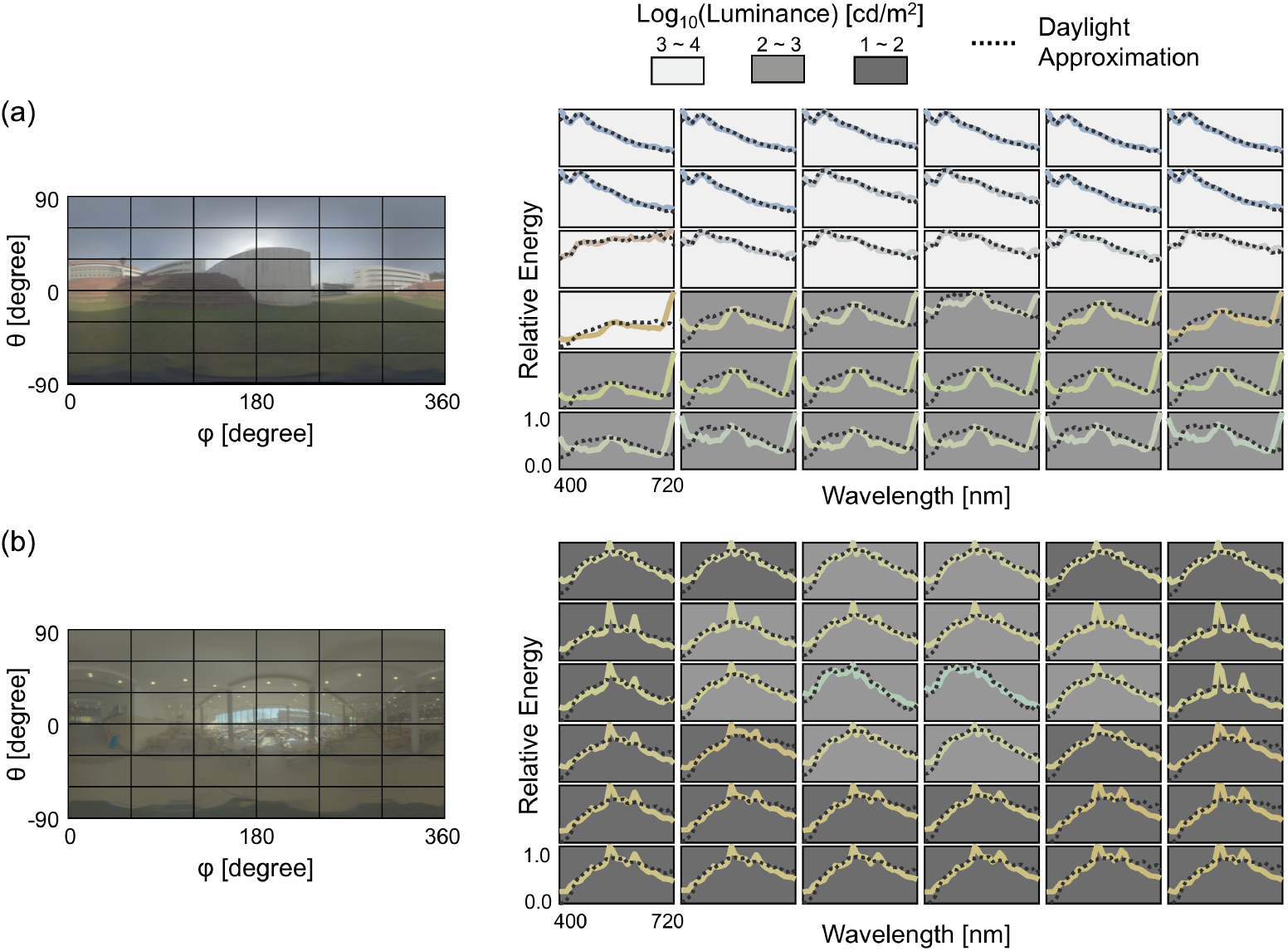
Directional spectral variation as a function of azimuth *φ* and elevation *θ*. (a) Example of one outdoor scene (campus). Each plot on the right-hand panel represents the mean spectrum within the corresponding rectangle drawn on the left-hand image. The colour of each curve is the sRGB colour of the spectrum. The background colour represents the log luminance level. The black dotted line indicates the spectrum derived from CIE daylight basis functions. (b) The case for one indoor scene (canteen). See Appendix for 10 other scenes.

Note that the black dotted curve in each plot represents the spectrum reconstructed using tristimulus values of the spectrum and the daylight-basis assumption (see equations (2a), (4a), (5a), [2]). Strictly speaking, these equations can be applied only to chromaticities on the daylight locus, but here we attempt to test the extent to which a linear weighted sum of daylight basis functions accounts for observed spectral variation. The reconstructed spectrum based on the daylight assumption was normalized so that it had an equal area under the curve as that of the measured spectrum. It is clear that the reconstructed spectra are not well matched to the real spectra for lower regions of the image, corresponding to low elevation angles. As expected, this demonstrates that not all natural illuminant spectra can be recovered from daylight basis functions in a scene where secondary reflection exists. For the canteen scene (b), spectral shapes are much more uniform across incident angle. Overall, spectral shapes rather resemble the CIE daylight function, presumably because the artificial overhead lighting from fluorescent tubes was mixed with daylight entering the room from the large window. It is notable that the regions oriented towards the large window have exceptionally high luminance levels.

It should be noted this trend depended on the scene, as shown in the Appendix, Figures S1 and S1. To see whether the similarity between observed spectra and reconstructed spectra by CIE daylight basis functions differs between the upper hemisphere (where directly emitted lights dominate) and the lower hemisphere (where the secondary reflection dominates), we calculated the root mean square error (RMSE) between the measured spectra and the reconstructed spectra for all panels in Figure 4. Note that in Figure 4, all spectra were scaled so that maximum value of the spectrum becomes one within each panel for the sake of visibility. However, when we calculate RMSE, all spectra used for calculation were scaled so that they have the same area under the curve. This normalization allows us to compare spectral shape whilst ignoring intensity differences. Due to this normalization procedure, the units of these RMSE values were arbitrary. The RMSE values were separately averaged across the upper 18 panels (theta >= 0) and the lower 18 panels (theta < 0) for each scene. Figure 5 shows the mean RMSE values across upper panels and lower panels for (a) 8 outdoor scenes and (b) 4 indoor scenes.

**Fig. 5.**
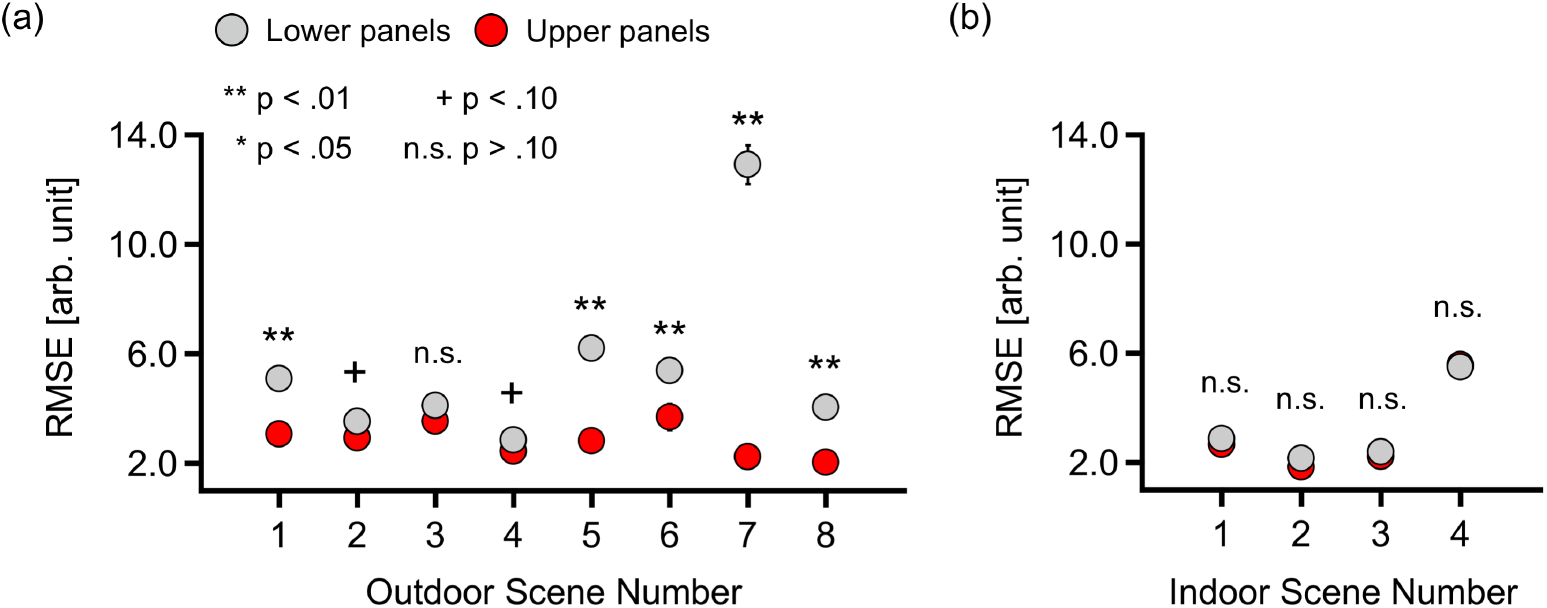
Average RMSE between measured spectra and reconstructed spectra based on the CIE daylight assumption, across the upper 18 panels (red circle symbols) and the lower 18 panels (gray circle symbols). The panels (a) and (b) show 8 outdoor scenes and 4 indoor scenes, respectively. The p-values to show whether the upper part and lower part have significantly different RMSE values are indicated by signs above each symbol. The error bars indicate ± S.E. but their lengths were mostly smaller than the symbols.

A two-way repeated-measures analysis of variance (ANOVA) was performed for (a) outdoor scenes with scene (scene 1 to scene 8), and hemisphere (upper and lower) as within-subject factors for the RMSE values. For (a) outdoor scenes, the main effect of hemisphere, scene and the interaction between two factors were all significant (F(1,17) = 512.5, *η*^2^ = 0.968, p < .00001; F(7,119) = 40.5, *η*^2^ = 0.704, p < .00001; F(7,119) = 52.6, *η*^2^ = 0.756, p < .00001). The analysis of interaction revealed that for scene 1, 5, 6, 7 and 8, the lower part showed significantly higher RMSE values (F(1,17) = 26.5, p = .000081; F(7,17) = 43.4, p < .00001; F(1,17) = 9.34, p = .007145; F(1,17) = 214.3, p < .00001; F(1,17) = 52.8, p < .00001, respectively). In contrast, scene 2, 3, 4 did not show significant difference between upper and lower hemisphere (F(1,17) = 3.68, p = .072; F(7,17) = 2.16, p = .160; F(1,17) = 3.38, p = .084, respectively). For (b) indoor scenes, the main effect of hemisphere was not significant (F(1,17) = 1.35, *η*^2^ = 0.282, p = .261). However, the main effect of scene was significant (F(3,51) = 73.41, *η*^2^ = 2.08, p < .00001). The interaction between the two factors was not significant (F(3,51) = 0.59, *η*^2^ = 0.186, p = .624). Thus, the dissimilarity of the measured spectra to that recovered from the daylight basis functions is similar between upper and lower hemispheres. Overall, for outside scenes, secondary reflection alters the spectrum shape substantially, and thus the daylight assumption does not adequately account for the observed spectral variation. In contrast, for indoor scenes, dissimilarity (RMSE) was small overall compared to outdoor scenes, and the difference between upper and lower hemispheres was essentially absent.

To compare the uniformity of spectral shape across direction, Figure 6 shows heat maps in which each pixel indicates the RMSE between the measured spectrum and the mean spectrum across all pixels in that scene. Values at the top left of each image indicate the averaged RMSE across all pixels. If the scene had the same spectra at every pixel, the heatmap would be uniformly black. Note that the mean spectrum across all pixels was calculated based on raw data, and thus it was dominated by intense spectra. However, again, when we calculated the RMSE, all spectra were normalized to have equal area so that we can compare the dissimilarity of spectral shape whilst ignoring intensity differences. Overall, the mean RMSE seems to be smaller for indoor scenes than for outdoor scenes. The mean of averaged RMSE values (shown at the top left in each panel) across 8 outdoor scenes was 7.95 and that for indoor scenes was 5.49, which was shown to be significantly different (p = 0.0121, t(10) = 3.06, t-test). Therefore, the indoor scenes have rather more uniform directional structure of spectral shapes across all directions than outdoor scenes.

**Fig. 6.**
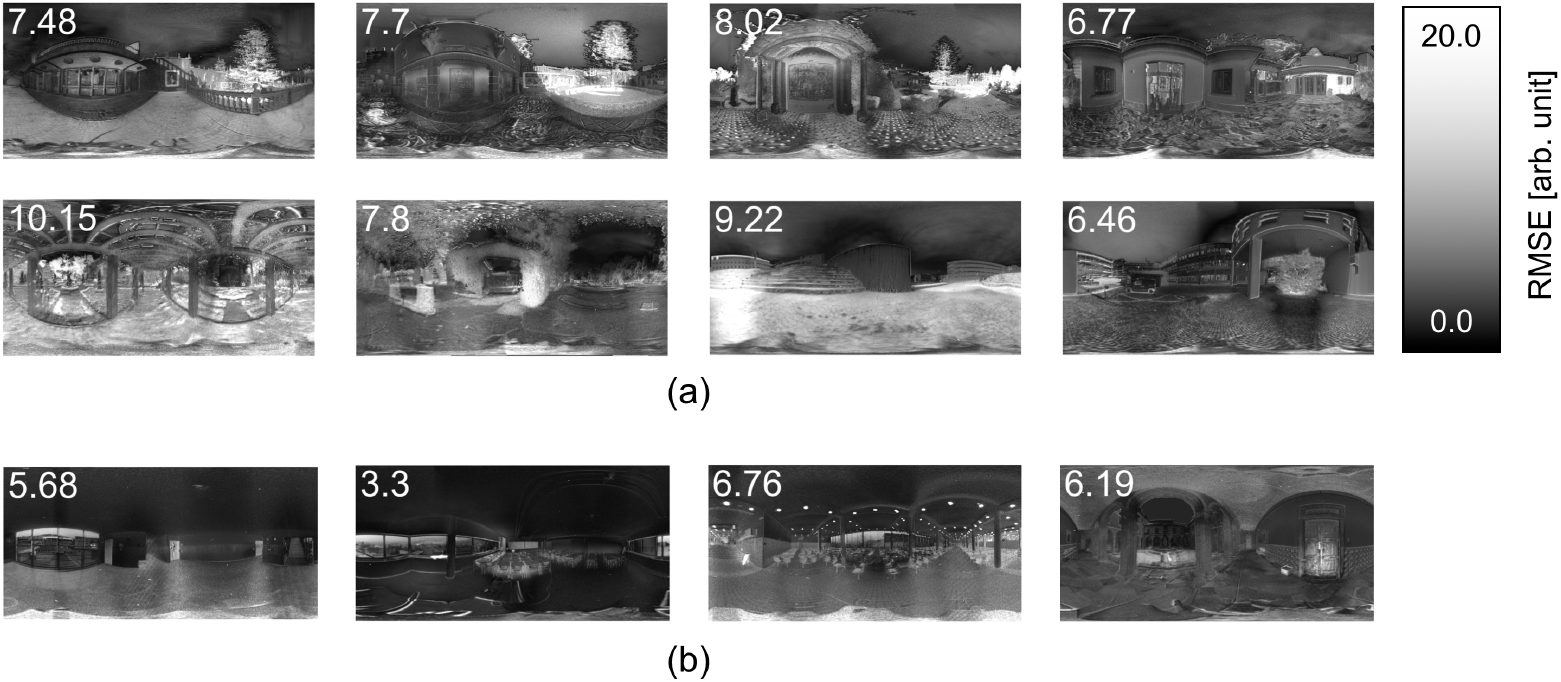
Heatmap to show the spectral variation for (a) outdoor scenes and (b) indoor scenes. Each pixel indicates the RMSE between measured spectrum and the mean spectrum across all pixels. The units are arbitrary. Note spectra in all pixels and the mean spectrum were normalized to have the equal area to allow us to compare the spectral shape. This calculation was performed independently for each scene so that the overall luminance differences within each scene did not affect the calculation. The number at the top left of each image indicates the mean RMSE value across all pixels.

Additionally, we characterize the spatial structure of these environmental illumination maps, to understand the variation in spatial scale from one scene to another, and also from one spectral channel to another. For 2D image planes, the power spectrum of the 2-D Fourier transformation would answer this question, but since environmental illumination maps are specified in spherical coordinates we used a spherical-harmonic decomposition. Figure 7 (a) shows a schematic illustration. The right-hand part of the panel shows the series of spectral images that comprise the original hyperspectral image (left-hand side). Spherical harmonics adopt two parameters, namely degree and order, that control the frequency and alignment of the basis functions. Detailed explanation is documented elsewhere (e.g. [23]). In panels (b) and (c) we show the power spectra (i.e. the absolute value of the spherical harmonic coefficient) as a function of degree *l*. Coefficients were summed across orders, and each curve was normalized independently so that the highest point within the curve becomes zero. Each point was connected by spline interpolation. For all scenes, the power is highest at zero or small degrees, and decreases rapidly as the degree increases. Scenes marked by plus at the bottom left in the panel shows a particularly strong wavelength dependency that the power of short wavelengths decrease more rapidly than that of longer wavelengths. Therefore, for some scenes longer wavelengths seem to have finer spatial structure. However, note that this wavelength dependency was not observed for all scenes.

**Fig. 7.**
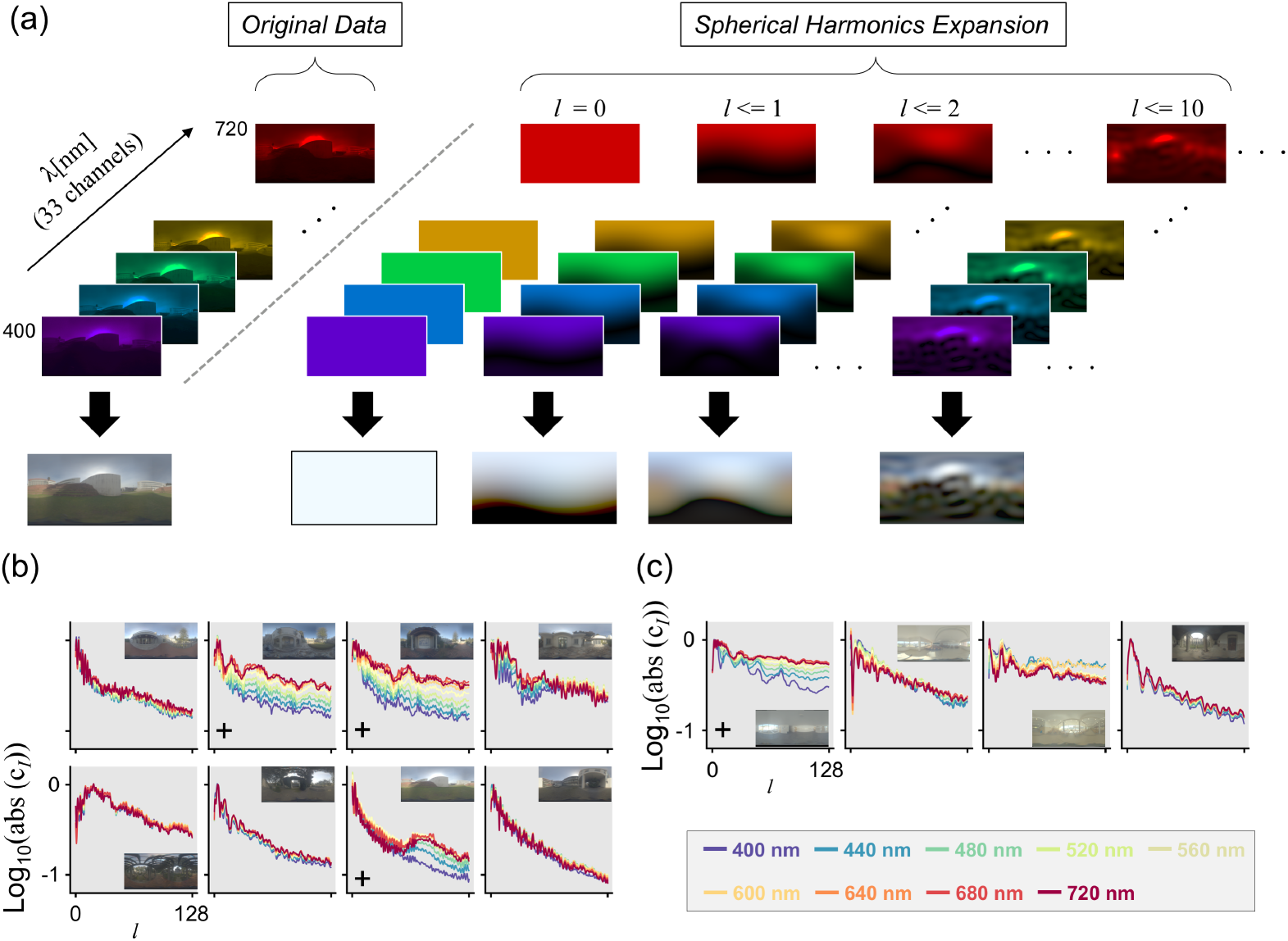
The concept of spherical-harmonic expansion. (a) Spherical-harmonic expansion was performed independently for each wavelength. The variable *l* represents the degree and the right-hand panel shows the reconstructed images up to a specified degree level. (b) Power spectra of spherical-harmonic expansion for wavelengths from 400 nm to 720 nm every 40 nm as a function of degree *l* for outdoor scenes. (c) Power spectra of indoor scenes.

One important implication from the present findings relates to human perceptual colour constancy, which is the ability of the human colour vision system to support the stable perception of an object’s surface colour regardless of illumination. Most previous studies of human colour constancy have used a simplified experimental setting where an object is illuminated by a single-spectrum illumination [24]. Under such a lighting environment, the computation to achieve colour constancy can be conceptualized as first estimating the illumination colour and then discounting its influence from the whole visual field. This approach is not plausible in real-world scenes because an object receives different lights from every direction. Moreover, by slightly changing head or object position, the object samples a different set of incident lights. Nevertheless, various research suggests that our perception of surface properties of objects such as shape, colour or gloss is not critically impaired under such complex lighting environments [25–34]. Another notable feature of the present data is the failure of the daylight assumption for lights from below. This is consistent with one study on colour constancy that suggests the visual system may not actually rely on the daylight assumption [35].

One limitation of the present study was the spatial resolution of the image due to the use of a mirror sphere and limited resolution of the hyperspectral camera itself (the image of the sphere was sampled at 1024 × 1024 pixels). In addition, the present measurements avoided direct sunlight, and thus the dynamic range was also limited. Hyperspectral measurement of a single sphere image can take over 10 minutes to capture sufficient light for all wavelengths. We prioritized the data acquisition time over the dynamic range, which would require a change of the exposure time, because otherwise the lighting environment is likely to change during the measurement. Methods to measure multispectral environmental illuminations have been improving over recent years [36, 37], and it will be important to measure additional hyperspectral datasets for further generalization of the findings. Nevertheless, the present study provided an important first step toward understanding the complex nature of environmental illumination, especially in the spectral domain.

## 4. Conclusion

Based on novel datasets collected in this study, we found that the chromaticites of natural illuminants are not restricted to the daylight locus, but instead show a much wider chromatic variation than previously thought. More importantly, the data clearly showed that natural lighting environments possess significant directional spectral variation. The degree of variation seems to be much larger in outdoor scenes than in indoor scenes.

## Acknowledgments

Authors acknowledge Museu Nogueira da Silva for the use of their garden to collect datasets and Andreia E. Gomes, Catarina F. M. Herdeiro, Eduardo G. Vicente and Joshua Harvey for the help of measurement. Authors also thank David H. Foster and Tom C. B. McLeish for advice on image processing.

## Supplemental Information

Figure S1 and S2 show a directional spectral variation for other outdoor and indoor scenes, respectively. Here, we see that the degree of spectral variation somewhat depends on the scene. However, we observe a similar general trend as described in the main text. That is, for outdoor scenes as shown in Figure S1, light from above tend to be more bluish and high luminance while lights from below have less energy around short-wavelength region and intensity is weaker. Indoor scenes have relatively uniform spectral shape regardless of direction, but region that has a large window or a region open to the sky have exceptionally high luminance.

