## Supplementary figures and images for "Hyperspectral environmental illumination maps: Characterizing directional spectral variation in natural environments"

### Supplemental Figure S1

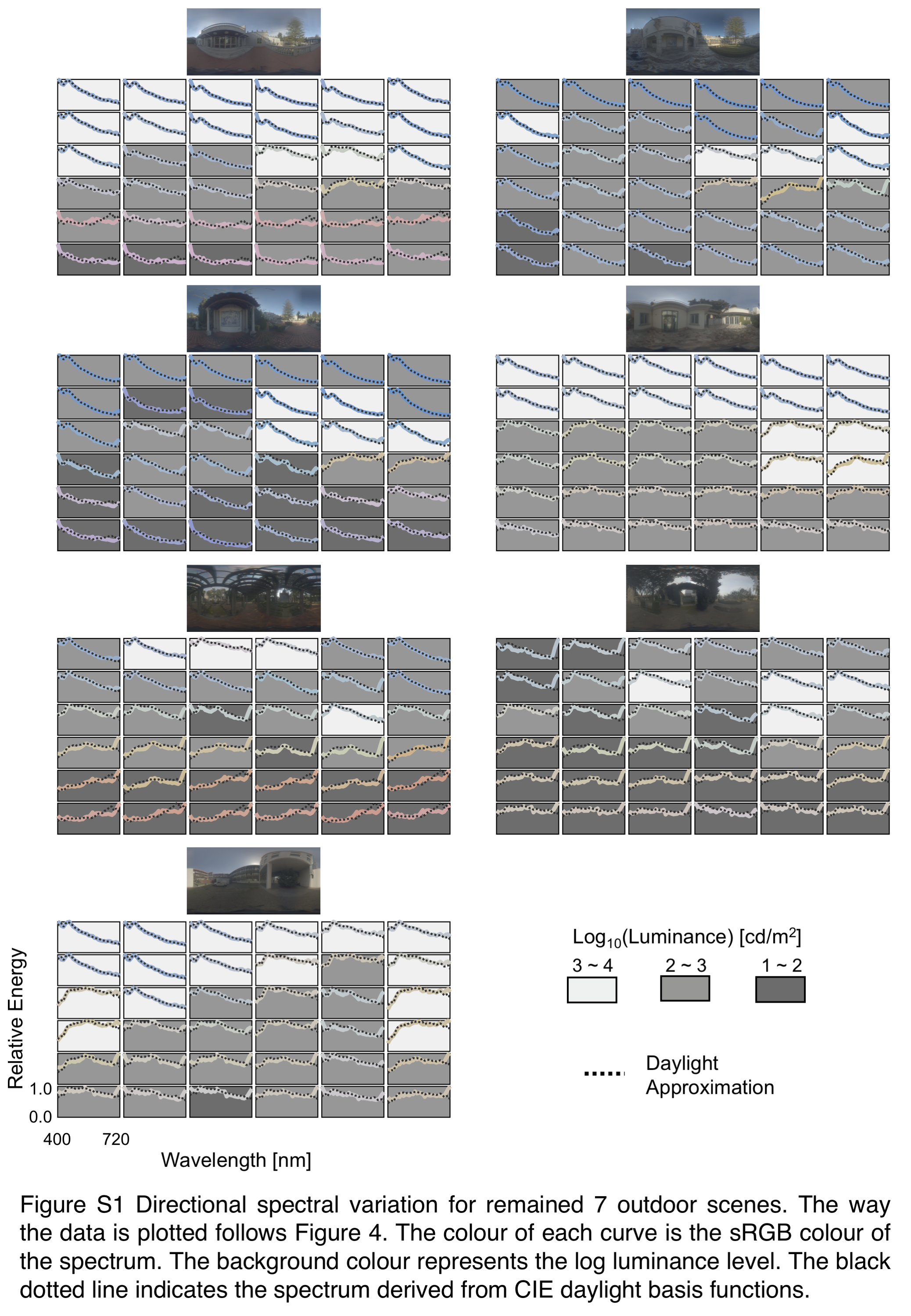

### Supplemental Figure S2

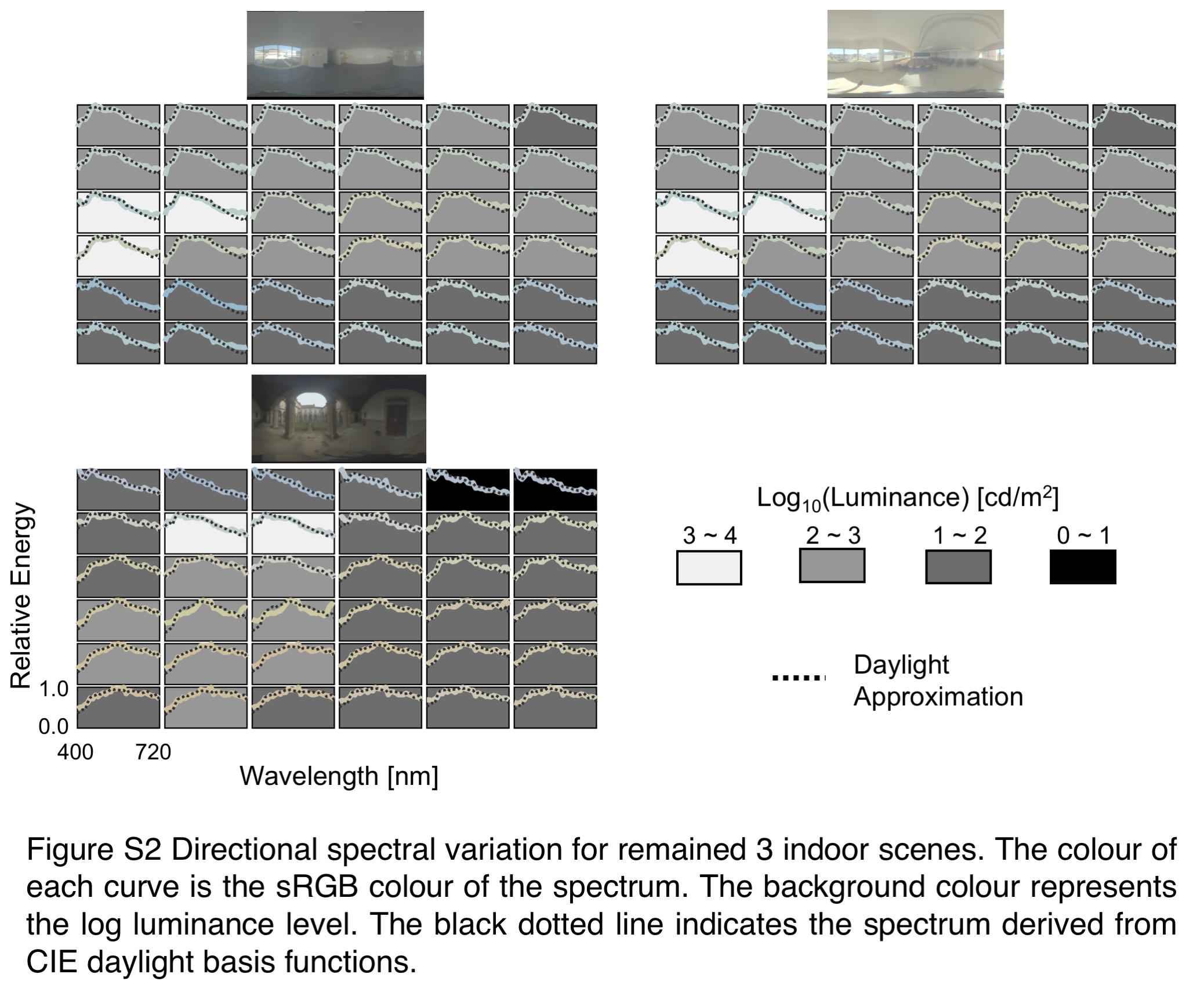
